# Regulation of plasmodesmata at specific cell-cell interfaces

**DOI:** 10.1101/2021.02.25.432931

**Authors:** Zhongpeng Li, Kyaw Aung

## Abstract

Precise exchange of information and resources among cells is essential for multicellular organisms. Intercellular communication among diverse cell types requires differential mechanisms to achieve the specific regulation. Despite the significance of intercellular communication, it is largely unknown how the communication between different cells is regulated. Here, we report that two members of plasmodesmata-located proteins modulate plasmodesmata at two distinct cell-cell interfaces.

## Main

Plasmodesmata (PD) are membrane-lined channels connecting adjoining plant cells, allowing the exchange of signals and resources (Lucas et al., 2009). The PD-dependent communication among cells is fundamental for developmental regulation and stress responses in plants (Sager and Lee, 2018). PD allow translocating photosynthetic products from mature leaves (source tissues) to non-photosynthetic parts of the plant (sink tissues; Comtet et al., 2017; Liesche and Patrick, 2017). Recent findings also highlighted the crucial role of PD in plant immunity against filamentous and bacterial pathogens (Lee et al., 2011; Faulkner et al., 2013; Cao et al., 2015; Aung et al., 2020; Tomczynska et al., 2020). The function of PD is controlled by the homeostasis of a plant polysaccharide, callose. Callose synthases (CalSs) and β-1,3 glucanases mediate the biosynthesis and degradation of callose, respectively, at the plasma membrane of PD (De Storme and Geelen, 2014). Callose deposition at PD suppresses the PD-dependent movement of molecules between adjoining cells, presumably by narrowing the PD aperture. Other than callose, plasmodesmata-located proteins (PDLPs) also play important roles in regulating the plasmodesmal function, but how PDLPs regulate PD function is unclear (Lee et al., 2011; Wang et al., 2020).

To better understand the function of PDLPs, we individually overexpressed all eight members of PDLP (PDLP1-8; Thomas et al., 2008) in Arabidopsis wild-type Col-0. The PDLPs were fused with His and Flag (HF) tag. The levels of protein expression in the transgenic plants were assessed by immunoblot analyses using a Flag antibody (Figure 1a and Supplemental Figure 1a). Here, we report the characterization of *p35S::PDLP5-HF* and *p35S::PDLP6-HF*. Consistent with a previous report (Lee et al., 2011), a stunted growth phenotype was observed for the *p35S::PDLP5-HF* lines compared to a vector control *p35S::HF-YFP*. We also observed a positive correlation between the expression level of PDLP5 and the growth phenotype (Supplemental Figure 1a-1b; Lee et al., 2011). In addition, we identified *p35S::PDLP6-HF* transgenic lines with more extreme developmental phenotypes, including a stunted growth and late flowering. Similar to PDLP5, transgenic plants with higher expression level of PDLP6 exhibit a more severe dwarfism in plant growth (Figure 1a-1b and Supplemental Figure 1c). We hypothesized that the overexpression of PDLP5 and PDLP6 affects the PD-dependent sugar translocation for the following reasons: (1) PD are required for symplastic movement of sugars between mesophyll cells in mature leaves (Schulz et al., 2015), (2) PD are involved in sugar loading within phloem in Arabidopsis (Liesche et al., 2017), and (3) Arabidopsis mutants compromised in sugar translocation exhibit a stunted growth phenotype (Srivastava et al., 2008; Chen et al., 2012).

**Figure 1.**
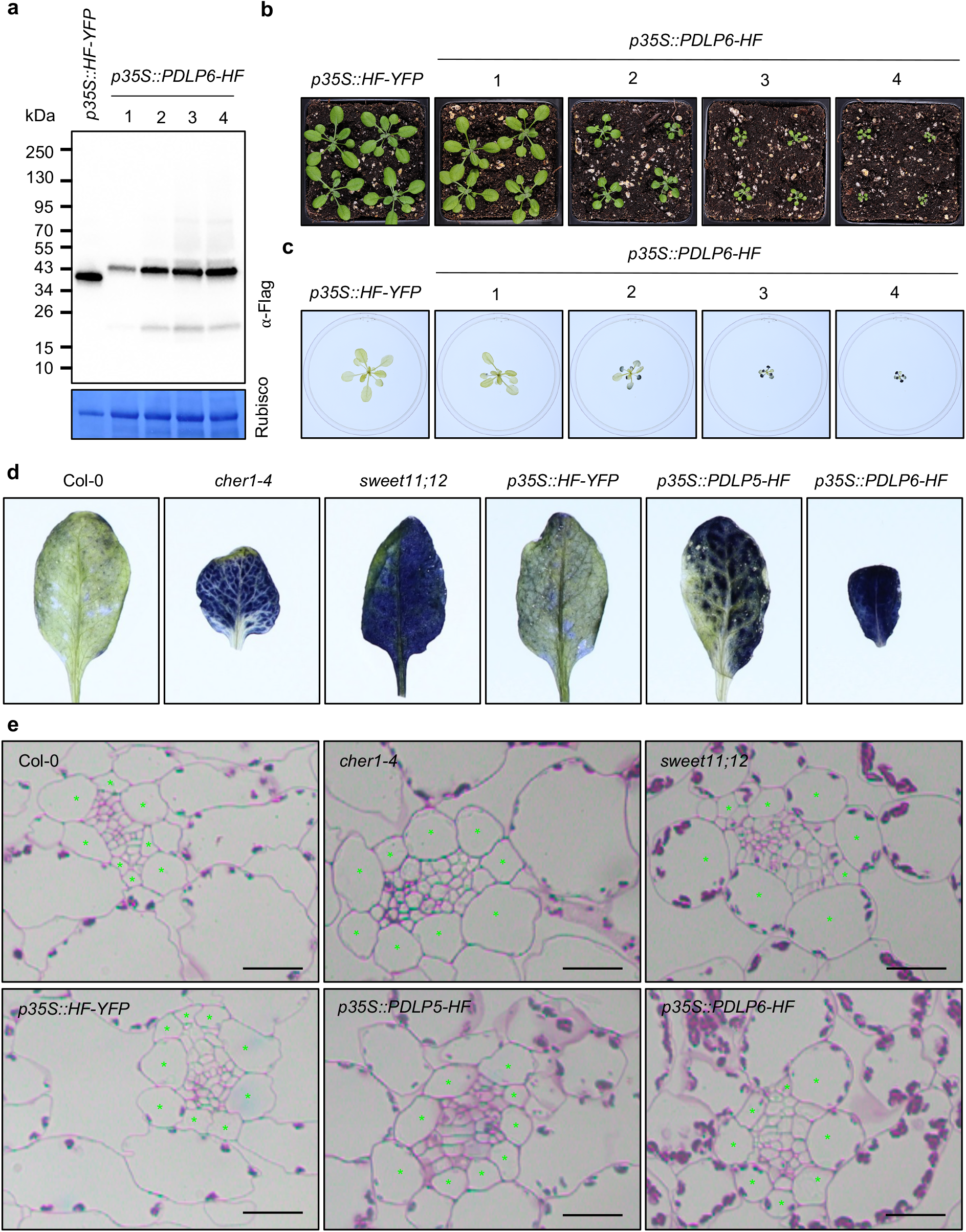
The overexpression of PDLP5 and PDLP6 leads to stunted growth and starch overaccumulation. **(a)** Immunoblot analysis detects the expression of PDLP6-HF in four independent transgenic plants. An anti-Flag antibody was used to detect the expression of Flag-fusion proteins. Rubisco was served as loading controls. **(b)** 3-week-old Arabidopsis plants were grown under a light intensity used for standard Arabidopsis growth (110 µmol m^-2^ s^-1^). **(c)** Starch accumulation phenotype of four independent transgenic lines expressing PDLP6-HF. The plants were grown under a light intensity used for standard Arabidopsis growth (110 µmol m^-2^ s^-1^) for three weeks. Plants were subjected for starch staining using Lugol’s solution at the end of the night. **(d)** Starch staining of 5-week-old Arabidopsis plants. 4-week-old Arabidopsis plants were irradiated with a high light intensity (200 µmol m^-2^ s^-1^) for a week. Samples were collected at the end of the night for starch staining using Lugol’s solution. All images were captured with the same magnification. **(e)** Histological sections of Arabidopsis leaves. Mature leaves of 5-week-old high light-treated plants were subjected to sectioning and staining with periodic acid/Schiff reagent, which stains polysaccharide in cell wall and starch grains in chloroplasts. Asterisks mark bundle sheath cells. Scale bars = 20 µm.

In mature leaves, chloroplasts within mesophyll cells (MCs) convert solar energy into chemical energy, producing sugars. The carbohydrate can be stored temporarily in the chloroplasts as starch or translocated to different parts of the plants. To translocate photosynthetic products, sugars move from MCs to sieve elements (SEs) through different pathways. It’s generally believed that photosynthetic products move symplastically through PD between photosynthetic cells, including MCs and bundle sheath cells (BSs), and are loaded into phloem for long-distance transport (Lucas et al., 2013). In Arabidopsis, three major cell types make up phloem: phloem parenchyma cells (PPs), companion cells (CCs), and SEs. Within phloem, symplastic and apoplastic phloem loading have been reported for different plant species. Symplastic phloem loading moves sugars mainly through PD connecting different phloem cell types. Arabidopsis utilizes apoplastic phloem loading, in which sugars are exported from PPs into the extracellular space and imported into CCs. From CCs, sugars move symplastically into SEs for long-distance transport. Arabidopsis SUGARS WILL EVENTUALLY BE EXPORTED TRANSPORTER (SWEET) proteins (SWEET11 and SWEET12) function as sugar exporters on the plasma membrane of PPs (Chen et al., 2012), whereas a sucrose-proton symporter 2 (SUC2) protein imports sugars from the apoplast into CCs in Arabidopsis (Srivastava et al., 2008). *sweet11;12* and *suc2* Arabidopsis mutants exhibit compromised sugar translocation from source to sink, resulting in overaccumulation of starch in mature leaves and stunted growth (Srivastava et al., 2008; Chen et al., 2012). Sugars from CCs move into SEs through PD (Lucas et al., 2013; Liesche and Patrick, 2017), completing the sugar-loading process. The enucleated SEs transport sugars to other parts of the plants (Oparka and Turgeon, 1999; Chen et al., 2015; Zhang and Turgeon, 2018). In a simplified pathway, sugars move from MCs→BSs→PPs→CCs→SEs in mature leaves to translocate sugars in Arabidopsis.

To investigate whether the stunted growth phenotypes of the transgenic plants are due to defects in symplastic translocation of sugar in mature leaves, we first examined starch accumulation in *p35S::PDLP6-HF* transgenic plants. The aerial portions of 3-week-old plants were stained with Lugol’s solution at the end of the night, when overall starch accumulation in mature leaves is at its lowest (Chen et al., 2012). When grown under a light intensity used for standard Arabidopsis growth (110 µmol m^-2^ s^-1^), 3 out of 4 *p35S::PDLP6-HF* independent transgenic plants overaccumulate starch in mature rosette leaves, exhibiting a dark blue color, compared to *p35S::HF-YFP*. Similar to plant growth phenotype, transgenic plants with higher expression level of PDLP6 contain more starch in their mature leaves (Figure 1c). For further analyses, we choose *p35S::PDLP6-HF* (#2). We also determined starch accumulation phenotype in *p35S::PDLP5-HF* (#1) due to its stunted growth phenotype (Supplemental Figure 1). To compare starch accumulation phenotype with mutants defective in sugar translocation, we included Arabidopsis mutants, *cher1-4* (Kraner et al., 2017) and *sweet11;12* (Chen et al., 2012) as both mutants overaccumulate starch in leaves. Consistent with previous report (Kraner et al., 2017), *cher1-4* shows starch overaccumulation in mature rosette leaves. Col-0, *p35S::HF-YFP, sweet11;12*, and *p35S::PDLP5-HF* all display comparable starch accumulation phenotype (Supplemental Figure 2a). As *sweet11;12* exhibits starch accumulation only when the plants were grown under a high light condition (Chen et al., 2012), we irradiated the mutants and transgenic plants with a high light intensity (200 µmol m^-2^ s^-1^) for a week. The starch overaccumulation phenotype became apparent in *sweet11;12* as well as *p35S::PDLP5-HF*, when the plants were grown under a high light condition (Figure 1c and Supplemental Figure 2b). Intriguingly, we observed distinct starch accumulation patterns among the genotypes. *p35S::PDLP6-HF* and *sweet11;12* accumulates starch evenly in mature leaves, whereas *p35S::PDLP5-HF* and *cher1-4* lack starch accumulation in and around vascular tissues in mature leaves (Figure 1d). As SWEET11 and SWEET12 specifically express at PPs within phloem (Chen et al., 2012), sugars are likely still able to move from MCs to PPs in the *sweet11;12* mutant. *cher1-4*, on the other hand, has a reduced number of PD between mesophyll cells to allow for symplastic movement of sugars (Kraner et al., 2017); consequently, it is possible that most sugars are trapped within the photosynthetic cells of the mutant. To determine cell type-specific starch accumulation in the mutants and transgenic plants, leaf discs from high light-treated plants were subjected to histological sectioning and starch staining. Periodic acid/Schiff (PAS) reagent labels polysaccharide in cell wall and starch grains in chloroplasts. Consistent with whole tissue staining (Figure 1d), *cher1-4, sweet11;12, p35S::PDLP5-HF*, and *p35S::PDLP6-HF* exhibit darker PAS-stained starch grains in chloroplasts of mesophyll cells compared to that of Col-0 and *p35S::HF-YFP*. In line with our prediction, *cher1-4* and *p35S::PDLP5-HF* contain fewer starch grains in BSs, whereas *sweet11;12* and *p35S::PDLP6-HF* contain many starch grains in their BSs. We also observed starch grains in PPs of *sweet11;12* (Figure 1e). Together, the findings support that the sugar translocation is likely blocked between (1) MC-MC and MC-BS in *cher1-4* and *p35S::PDLP5-HF*, (2) BS-PP in *p35S::PDLP6-HF*, and (3) PP-CC in *sweet11;12*. The distinct patterns of starch accumulation between *p35S::PDLP5-HF* and *p35S::PDLP6-HF* suggest that PDLP5 and PDLP6 express in and function at specific cell types to regulate the plasmodesmal function at two distinct cell-cell interfaces.

To determine whether PDLP5 and PDLP6 express in a cell-type-specific manner, we generated native promoter driven *pPDLP5::PDLP5-YFP* and *pPDLP6::PDLP6-YFP* transgenic Arabidopsis plants. T_3_ generations of the transgenic plants were subjected to confocal imaging. The expression of PDLP5-YFP was mainly detected between epidermal cells, whereas no PDLP5-YFP signals were observed in mesophyll cells and leaf vasculature. PDLP6-YFP, on the other hand, was expressed specifically in leaf vasculature (Figure 2a). We postulate that PDLP6-YFP expresses exclusively in PP as *PDLP6* transcripts were detected predominantly in the cell type (Kim et al., 2021). Similarly, PDLP6-YFP expresses specifically in root vasculature, whereas PDLP5-YFP was detected in epidermis and cortex (Figure 2a). Transgenic plants co-expressing PDLP-YFP and cell type-specific markers tagged with a compatible fluorescent protein (e.g., red fluorescent protein) for simultaneous imaging will further confirm the cell type-specific expression of PDLP6 in phloem. In parallel, we determined the promoter activity of *PDLP5* and *PDLP6* using β-Glucuronidase (GUS) reporter gene assay. Two transgenic lines, *pPDLP5::GUS* and *pPDLP6::GUS*, were subjected to GUS activity staining as previously described (Li et al., 2016). Among 20 independent *pPDLP5::GUS* transgenic plants, 15 of them exhibit a much higher GUS activity in leaf epidermal cells. The majority (13 out of 19) of *pPDLP6::GUS* transgenic plants displays the highest activity in leaf vasculature (Supplemental Figure 3). Representative transgenic lines of *pPDLP5::GUS* and *pPDLP6::GUS* were selected to analyze GUS activity in roots. *pPDLP5::GUS* transgenic plants exhibit GUS activity in epidermis and cortex, but not in the vasculature. *pPDLP6::GUS* transgenic plants, on the other hand, exhibit the highest GUS activity in the vasculature, likely phloem (Supplemental Figure 3). Together, our findings demonstrate that PDLP5 and PDLP6 are expressed in distinct and non-overlapping cell types. Interestingly, PDLP5 was reported to express in lateral root primordium-overlaying cells during lateral root emergence process (Sager et al., 2020). Further investigation will reveal the cell type-specific expression and function of PDLPs at different developmental stages and tissues.

**Figure 2.**
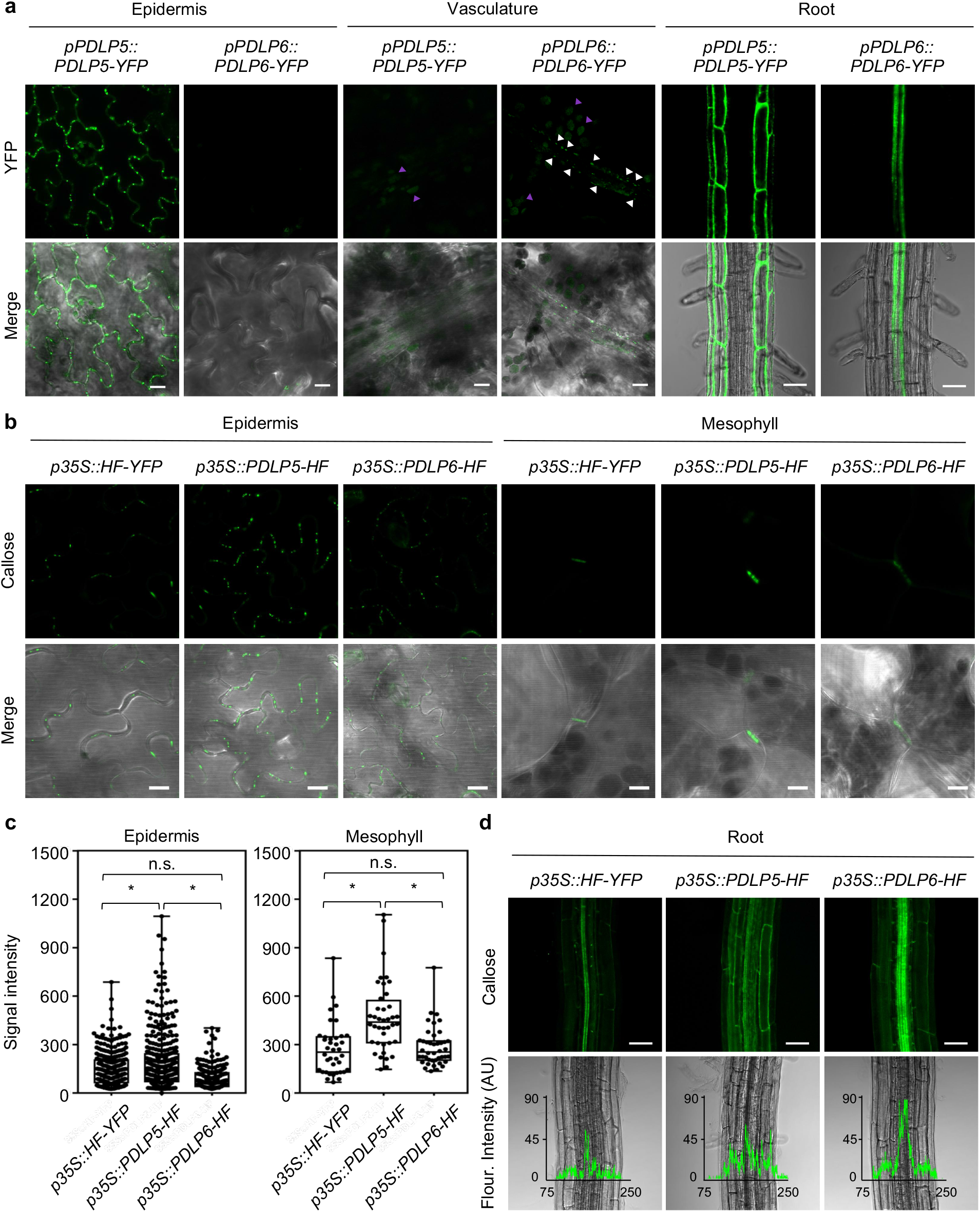
PDLP5 and PDLP6 express in and function at different cell types. **(a)** The cell type-specific expression of PDLP5-YFP and PDLP6-YFP. The fusion proteins were driven by their respective native promoter. Confocal images were captured from 2-week-old Arabidopsis seedlings. Green signals indicate the expression of the YFP fusion proteins in different cell types (top panel). Merged images of the signals from YFP and bright field are shown in the lower panel. White arrow heads indicate the expression of PDLP6-YFP at PD of vascular cells. Magenta arrow heads indicate chlorophyll autofluorescence. Scale bars for epidermis and vasculature = 10 µm. Scale bars for root = 50 µm. **(b)** Callose accumulation between leaf epidermal or mesophyll cells in transgenic plants overexpressing HF-YFP (control), PDLP5-HF, or PDLP6-HF. Green signals present aniline blue-stained callose at PD (top panel). Merged images of the signals from callose and bright field are shown in the lower panel. Scale bars = 10 µm. **(c)** Quantitative data show callose accumulation between epidermal cells and mesophyll cells. Asterisks indicate differences that are statistically significant (Mann-Whitney *U* Test; p<0.0002). n.s.: not significant. **(d)** Callose accumulation in roots of the transgenic plants. Green signals present callose accumulation in different root cell types (top panel). Scale bars = 50 µm. Merged images of bright field and fluorescence intensity profile (arbitrary unit: AU) of callose accumulation along the horizontal lines are shown in the lower panel. Numbers on the X axis indicate distance (µm) across the region analyzed.

While the molecular mechanism underlying PDLP function is unclear, the expression level of PDLP5 is highly associated with callose accumulation at PD in plants (Lee et al., 2011; Li et al., 2020). Similarly, the expression of PDLP1 is required for callose accumulation at haustoria during downy mildew infection in Arabidopsis (Caillaud et al., 2014). To further investigate the cell type-specific roles of PDLP5 and PDLP6, we examined whether the overexpression of PDLP5 or PDLP6 leads to the overaccumulation of callose at specific cell-cell interfaces. To detect callose accumulation, *p35S::PDLP5-HF, p35S::PDLP6-HF*, and *p35S::HF-YFP* leaves were stained with aniline blue. Consistent with the previous report (Lee et al., 2011), we observed a higher accumulation of callose between epidermal cells in *p35S::PDLP5-HF* as compared to both *p35S::PDLP6-HF* and *p35S::HF-YFP*. We also detected a higher callose level at PD connecting mesophyll cells in *p35S::PDLP5-HF* (Figure 2b-2c). As PDLP6 expresses specifically in vasculature, we also examined the callose accumulation between vascular cells. To this end, roots of the transgenic plants were stained with aniline blue and imaged with confocal microscopy due to their tissue transparency. We observed a much higher accumulation of callose in epidermis and cortex of *p35S::PDLP5-HF* than *p35S::HF-YFP* and *p35S::PDLP6-HF*. In contrast, *p35S::PDLP6-HF* exhibits the highest accumulation of callose in vascular cells (Figure 2d). Fluorescence intensity profiles were consistent among data collected from 25 individual transgenic plants for each genotype (Supplemental Figure 4). These findings support that the overexpression of PDLP5 and PDLP6 leads to the higher accumulation of callose at specific cell-cell interfaces.

We observed the cell type-specific functions of PDLP5 and PDLP6 while examining the transgenic plants overexpressing *PDLP5* or *PDLP6* genes. When driven by the same *35S* constitutive promoter, PDLP5-YFP and PDLP6-YFP are expressed in most cell types (Supplemental Figure 5; Aung et al., 2020), suggesting that PDLP5-HF and PDLP6-HF express ubiquitously in the transgenic plants. From the starch and callose accumulation phenotypes, it is clear that the *35S* driven expression of PDLP6 alters the plasmodesmal function only in the cell types where it is natively expressed. This suggests that different members of PDLPs regulate PD in a cell type-specific manner. One possibility for why constitutive expression of PDLP6 does not lead to PD defects in all cell types is that PDLPs might require other cell type-specific proteins to regulate PD. This hypothesis is supported by the finding that PDLP5 functions together with CalS8 to maintain basal callose homeostasis at PD, whereas it works together with CalS1 to deposit callose at PD during salicylic acid-dependent plasmodesmata regulation in Arabidopsis (Cui and Lee, 2016). The findings imply that different CalSs might function together with different PDLPs to regulate callose accumulation at PD in specific cell types. Alternatively, other unknown cell type-specific partners or factors might contribute to the PDLPs-dependent function in unique cell types. Although PDLP5-YFP is not detected in mesophyll cells when driven under the native promoter, the overexpression of PDLP5-HF leads to callose overaccumulation between mesophyll cells (Figure 2b-2c) and likely blocks sugars movement between photosynthetic cells. It’s plausible that the expression of PDLP5-YFP is under detectable threshold using confocal microscopy. Alternatively, functional partners of PDLP5 might be present in mesophyll cells, modulating callose accumulation and the plasmodesmal function.

The findings in this article begin to reveal how communication among distinct cell types is regulated in multicellular organisms. Further research should uncover the function of PDLPs and their partner proteins in regulating PD at the molecular level. As PD are critical in plant growth and defense, understanding the cell type-specific regulation of PD will provide a valuable toolkit to improve crop production.

## Methods

### Plant Material, Growth Conditions, Transformation, and Plant Selection

Arabidopsis (*Arabidopsis thaliana*) plants were grown at 22°C with 50% humidity and irradiated with 110 µmol m^-2^ s^-1^ white light for 14 h per day. To grow plants under a higher light intensity, plants were irradiated with 200 µmol m^-2^ s^-1^ white light. Transgenic Arabidopsis plants were generated using the simplified transformation method (https://plantpath.wisc.edu/simplified-arabidopsis-transformation-protocol/). To select for transgenic plants harboring transgenes containing the hygromycin-resistance gene, T_1_seeds were selected on 0.5× LS medium containing 25 μg/mL hygromycin. For those harboring transgenes containing the glufosinate-resistance gene, T_1_seeds were germinated on soil and 1-week-old seedlings were sprayed with 0.1% (v/v) Finale Herbicide (Bayer) and 0.05% (v/v) Silwet L-77 (PhytoTech). The T_2_or T_3_ plants were selected on 0.5× LS medium containing 10 μg/mL glufosinate-ammonium.

### Gene Cloning and Plasmid Construction

The coding sequences of PDLP1-8 were amplified with Gateway-compatible primers from cDNA synthesized from total RNA extracted from wild-type (Col-0) seedlings, and the coding sequence of yellow fluorescent protein (YFP) from pEarleyGate 101 (Earley et al., 2006) using Phusion High-Fidelity DNA polymerase (ThermoFisher). To generate *p35S::PDLP-HF* or *p35S::HF-YFP*, the PCR fragments were cloned into the pDONR 207 entry vector, followed by the pB7-HFC or pB7-HFN destination vector (Lee et al., 2018) using a standard Gateway cloning system (Invitrogen). To generate *pPDLP5::PDLP5-YFP* and *pPDLP6::PDLP6-YFP* constructs, a 3029 and 3331 bp including the promoter and PDLP genomic fragment with all exons and introns without stop codons were amplified from genomic DNA of wild-type Col-0 using Gateway-compatible primers. The PCR fragments were cloned into the pDONR 207 entry vector, followed by the pGWB540 destination vector (Nakagawa et al., 2007) using a standard Gateway cloning system (Invitrogen). To generate *pPDLP5::GUS* and *pPDLP6::GUS* constructs, the promoters (2051 bp and 1505 bp for PDLP5 and PDLP6, respectively) were amplified from genomic DNA of wild-type Col-0 using Gateway-compatible primers. The PCR fragments were cloned into the pDONR 207 entry vector, followed by the pMDC163 destination vector (Curtis and Grossniklaus, 2003) using a standard Gateway cloning system (Invitrogen). All primers used for cloning are listed in Supplemental Table 1.

### Whole Tissue Starch Staining

The aerial portion of 4-week-old Arabidopsis plants grown under regular light were harvested at the end of the night. For high light-treated plants, 4-week-old Arabidopsis plants grown under regular light were moved into a growth chamber with the same growth conditions except a higher light intensity. The inflorescence was removed and the rosette leaves were decolored overnight with 95% ethanol. The samples were then washed with ddH_2_O and stained with Lugol’s solution (Sigma-Aldrich) for 10 minutes and rinsed with ddH_2_O. The images were captured with a Canon camera 2 hours after staining.

### Tissue Sectioning and Starch Staining

Leaf punch samples were collected and fixed in FAA fixative (5% formaldehyde, 5% glacial acetic acid, 50% ethyl alcohol). Samples were dehydrated through graded ethanol series (70, 85, 95, 100%) for 3-6 hours each concentration. Samples were infiltrated into LR White hard grade resin (Electron Microscopy Sciences) and polymerized at 55°C for 48 hours. Sections were made using a Leica UC6 ultramicrotome at 1.5 µm thickness. Sections were stained for non-soluble polysaccharides as the following: slides with sections were immersed in periodic acid for 5 minutes, rinsed in distilled water for 5 minutes, stained in Schiff’s reagent (Electron Microscopy Sciences) for 10 minutes, rinsed in running tap water for 5 minutes, and air dried. Dry slides were coverslipped using Permount mounting media (Fisher Scientific).

### Aniline Blue Staining

Mature leaves from 4-week-old Arabidopsis leaves were infiltrated with 0.1 mg/ml aniline blue in 1x PBS buffer (pH 7.4). Samples were imaged at 5 minutes after the dye infiltration using confocal microscopy. Ten-day-old seedlings were vacuum-infiltrated with 0.1 mg/ml of aniline blue in 1x PBS buffer (pH 7.4). Stained root tissues were imaged using confocal microscopy. Callose in mature leaves was quantified using FIJI. For epidermis, images were converted from lsm to tiff. 8-bit images were used for analysis. Black and white images highlighting callose were created by Auto Threshold which was set by RenyiEntropy white method. Particle Analysis tool was used to outline callose with size from 0.10 to 20 µm^2^ and circularity from 0.15 to 1.00. Quantitative numerical values in µm^2^ were then exported. For mesophyll cells, aniline blue-stained callose area was manually selected and the signal intensity was determined by measuring integrated density. Semi-quantitative evaluation of the relative level of aniline blue-stained callose in the root tissues of the transgenic plants were performed using FIJI. Images were converted from lsm to tiff. 16 bit images were used for analysis. Horizontal lines were drawn near the bottom of the images crossing all root cell types as shown in Figure 2d. Plot profile was used to generate a two-dimensional graph. Value on y-axis represents the relative signal intensity as arbitrary unit (AU).

### GUS Activity Staining

GUS staining was performed as previously described (Li et al., 2016) with minor modifications. Mature leaves and small seedlings were immersed in GUS solution (100 mM sodium phosphate buffer [pH 7.0], 10 mM Na_2_EDTA, 1 mM K_3_ [Fe(CN)_6_], 1 mM K_4_ [Fe(CN)_6_], 0.1% Triton X-100, and 1 mM X-Gluc). Samples were vacuumed for 10-40 min, followed by incubation in darkness at 37°C for 2-16 h. After staining, samples were de-stained in 75% ethanol. Images were taken using ZEISS Axio Observer.

### Confocal Imaging

All confocal images were captured with a confocal laser-scanning microscope (Zeiss LSM 700). A small piece of tissue was mounted with water on a glass slide. For leaf tissues, the abaxial side was imaged. YFP was excited at 514 nm and emission was collected over the range of 510–550 nm using SP555. Aniline blue-stained callose was excited at 405 nm and emission was collected over the range of 420–480 nm using SP555.

### Immunoblot Analyses

Arabidopsis leaves were frozen with liquid nitrogen and homogenized with 1600 miniG (SPEX). Protein extraction buffer (60 mM Tris-HCl [pH 8.8], 2% [v/v] glycerol, 0.13 mM EDTA [pH 8.0], and 1× protease inhibitor cocktail complete from Roche) was added to the homogenized tissues (100 µl/10 mg). The samples were vortexed for 30 s, heated at 70°C for 10 min, and centrifuged at 13,000*g* for 5 min at room temperature. The supernatants were then transferred to new tubes. For SDS-PAGE analysis, 10 µl of the extract in 1x Laemmli sample buffer (Bio-Rad) was separated on 4–15% Mini-PROTEAN TGX precast protein gel (Bio-Rad). The separated proteins were transferred to a polyvinylidene fluoride membrane (Bio-Rad) using a Trans-Blot Turbo Transfer System RTA transfer kit following the manufacturer’s instructions (Bio-Rad). The membrane was incubated in a blocking buffer (3% [v/v] BSA, 50 mM Tris base, 150 mM NaCl, 0.05% [v/v] Tween 20 [pH 8.0]) at room temperature for 1 h, then incubated overnight with a 1:10,000 dilution of an α-Flag-HRP antibody (Sigma-Aldrich catalog No. A8592) at 4°C. The membrane was washed four times with 1× TBST (50 mM Tris base, 150 mM NaCl, 0.05% [v/v] Tween 20 [pH 8.0]) for 10 min. The signals were visualized with SuperSignal West Dura Extended Duration Substrate (Pierce Biotechnology).

## Acknowledgements

We thank the ABRC for providing the T-DNA insertion mutants and pMDC163 vector. We thank Tracey Stewart from Roy J. Carver high resolution microscopy facility at Iowa State University (ISU) for helping with histological sectioning and starch staining. We thank the Aung lab members Dr. Yani Chen, Dr. Su-Ling Liu, and Haris Variz. We also thank Dr. Yanhai Yin from ISU, Dr. Michelle Guo from ISU, Dr. Anne Rea from Michigan State University and Dr. Yu-Ti Cheng from Duke university for critically reading the manuscript. This work was supported by the National institute of General Medical Science (Grant R00GM115766 to K.A.).

## Contributions

K.A. and Z.P.L. designed the research. Z.P.L. conducted most experiments except the generation of *p35S::PDLP-HF* and *p35S::PDLP-YFP* transgenic lines. K.A. wrote the manuscript with inputs from Z.P.L.

## Figure legend

**Supplemental Figure 1. Characterization of *p35S::PDLP5-HF* and *p35S::PDLP6-HF* transgenic plants. (a)** Immunoblot analysis detects the expression of PDLP5-HF in two independent transgenic plants. An anti-Flag antibody was used to detect the expression of Flag-fusion proteins. Rubisco was served as loading controls. **(b)** Starch accumulation phenotype of two independent transgenic lines expressing PDLP5-HF. The plants were grown under a light intensity used for standard Arabidopsis growth (110 µmol m^-2^ s^-1^) for four weeks. Plants were subjected for starch staining using Lugol’s solution at the end of the night. **(c)** Stunted growth and late flowering phenotypes of Arabidopsis transgenic plants expressing *PDLP5-HF* or *PDLP6-HF. p35S::HF-YFP* was served as a control. Images were taken from 5-week-old plants using the same magnification.

**Supplemental Figure 2. Starch accumulation phenotype of Arabidopsis mutants and transgenic plants. (a)** The plants were grown under a light intensity used for standard Arabidopsis growth (110 µmol m^-2^ s^-1^; top panel) for four weeks. **(b)** For high light treatment, 4-week-old Arabidopsis plants were irradiated with a high light intensity (200 µmol m^-2^ s^-1^; lower panel) for a week. Samples were collected at the end of the night for starch staining using Lugol’s solution.

**Supplemental Figure 3. The promoter activities of Arabidopsis *PDLP5* and *PDLP6* genes.** Histochemical GUS analysis of Arabidopsis transgenic plants expressing the reporter genes under the control of the *PDLP5* or *PDLP6* gene promoter of Arabidopsis. Leaves of 4-week-old Arabidopsis T_1_ transgenic plants were subjected to GUS staining. Scale bar = 100 µm. Roots of 2-week-old Arabidopsis T_2_ transgenic plants were subjected to GUS staining. Scale bar = 50 µm. The numbers indicate the transgenic plants exhibit the shown GUS activity pattern out of the total independent transgenic plants analyzed.

**Supplemental Figure 4. Callose accumulation in roots of Arabidopsis transgenic plants overexpressing HF-YFP, PDLP5-HF, or PDLP6-HF**. Semi-quantitative evaluation of the relative level of aniline blue-stained callose in root cells was performed by analyzing the signal intensity across different root cell types. Confocal images were captured from the maturation zone of 10-day old seedlings. 25 individual transgenic plants were analyzed for each genotype and fluorescence intensity profiles (arbitrary unit: AU) of callose accumulation were combined within the genotype. Numbers on the X axis indicate distance (µm) across the region analyzed.

**Supplemental Figure 5. Ubiquitous expression of PDLP5-YFP and PDLP6-YFP fusion proteins in *p35S::PDLP5-YFP* and *p35S::PDLP6-YFP* transgenic plants.** The expression of the fusion proteins was detected in epidermal cells, mesophyll cells, vascular cells in leaves, and most cell types in roots. Confocal images were captured from 2-week-old Arabidopsis seedlings. Scale bars for epidermis, mesophyll, and vasculature = 10 µm. Scale bars for root = 50 µm.

**Supplemental Table 1. Primers used for cloning in this study.**

