## Supplemental Figures for "Regulation of plasmodesmata at specific cell-cell interfaces"

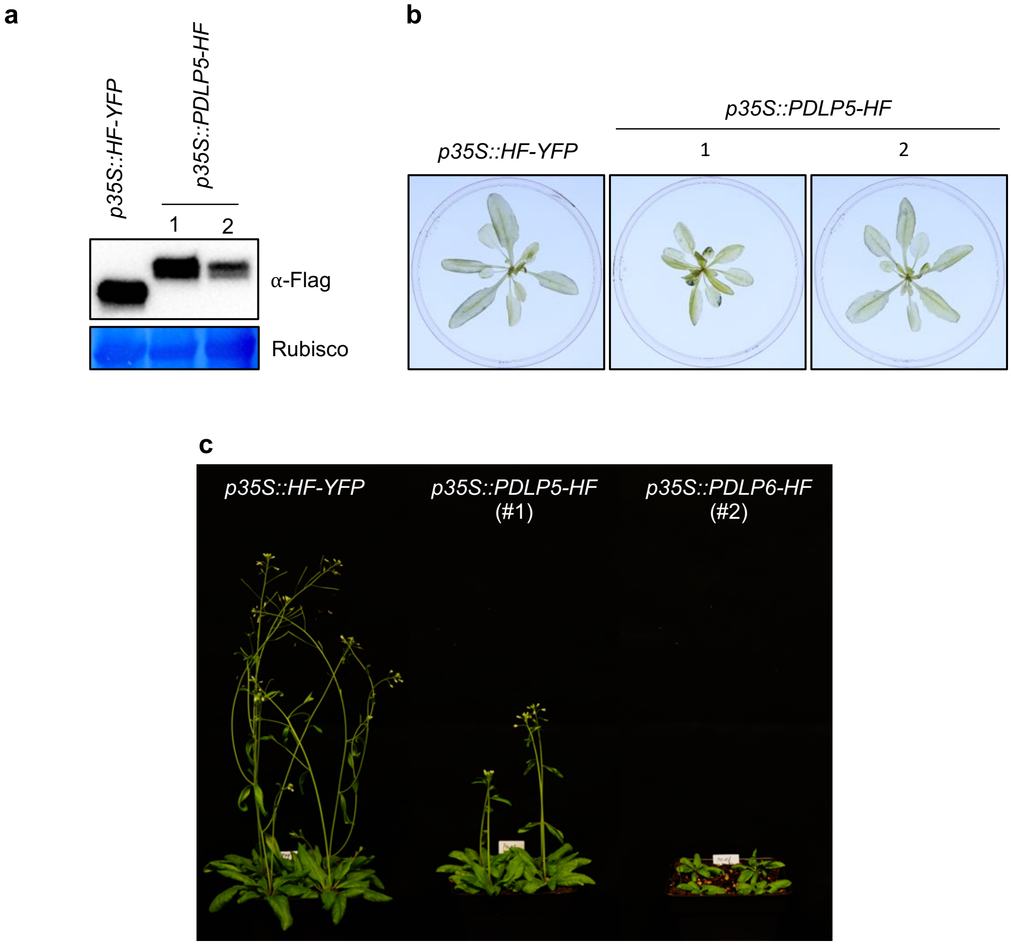


**Supplemental Figure 1. Characterization of *p35S::PDLP5-HF* and *p35S::PDLP6-HF* transgenic plants. (a)** Immunoblot analysis detects the expression of PDLP5-HF in two independent transgenic plants. An anti-Flag antibody was used to detect the expression of Flag-fusion proteins. Rubisco was served as loading controls. **(b)** Starch accumulation phenotype of two independent transgenic lines expressing PDLP5-HF. The plants were grown under a light intensity used for standard Arabidopsis growth (110 μmol $m^{-2}$ $s^{-1}$) for four weeks. Plants were subjected for starch staining using Lugol's solution at the end of the night. **(c)** Stunted growth and late flowering phenotypes of Arabidopsis transgenic plants expressing *PDLP5-HF* or *PDLP6-HF*. *p35S::HF-YFP* was served as a control. Images were taken from 5-week-old plants using the same magnification.

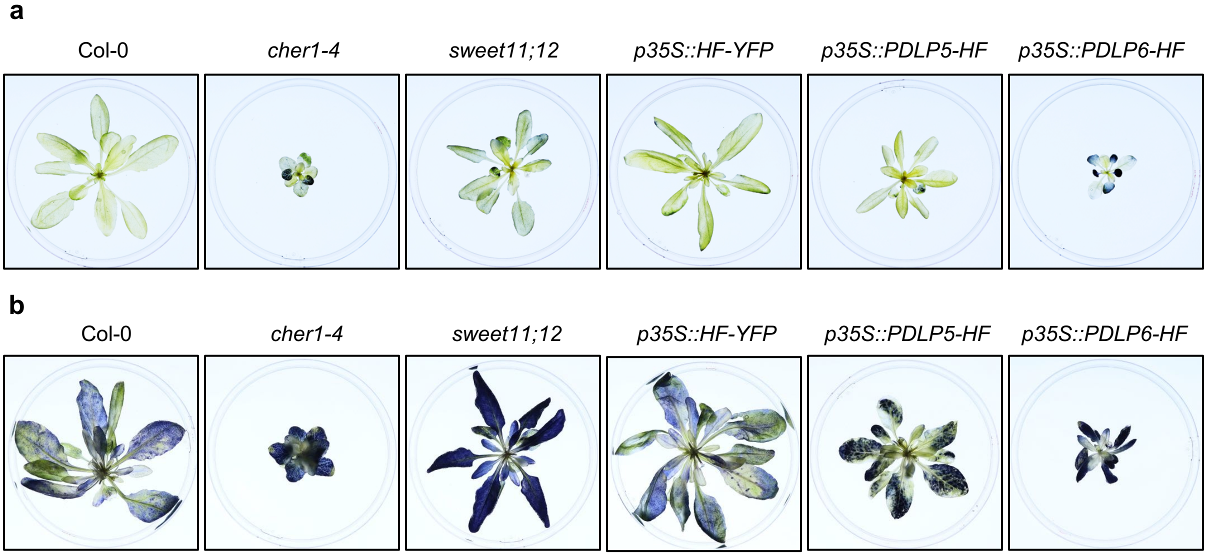


**Supplemental Figure 2. Starch accumulation phenotype of Arabidopsis mutants and transgenic plants. (a)** The plants were grown under a light intensity used for standard Arabidopsis growth (110 μmol $m^{-2}$ $s^{-1}$; top panel) for four weeks. **(b)** For high light treatment, 4-week-old Arabidopsis plants were irradiated with a high light intensity (200 μmol $m^{-2}$ $s^{-1}$; lower panel) for a week. Samples were collected at the end of the night for starch staining using Lugol's solution.

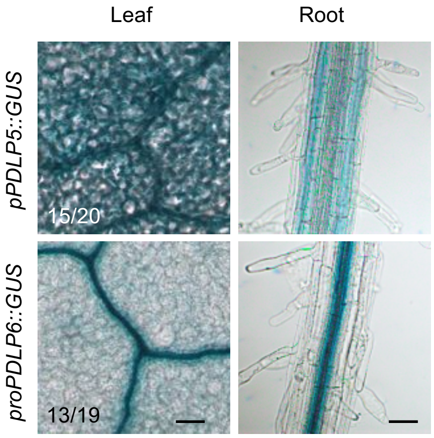


**Supplemental Figure 3. The promoter activities of Arabidopsis *PDLP5* and *PDLP6* genes.** Histochemical GUS analysis of Arabidopsis transgenic plants expressing the reporter genes under the control of the *PDLP5* or *PDLP6* gene promoter of Arabidopsis. Leaves of 4-week-old Arabidopsis $T_1$ transgenic plants were subjected to GUS staining. Scale bar = 100 μm. Roots of 2-week-old Arabidopsis $T_2$ transgenic plants were subjected to GUS staining. Scale bar = 50 μm. The numbers indicate the transgenic plants exhibit the shown GUS activity pattern out of the total independent transgenic plants analyzed.

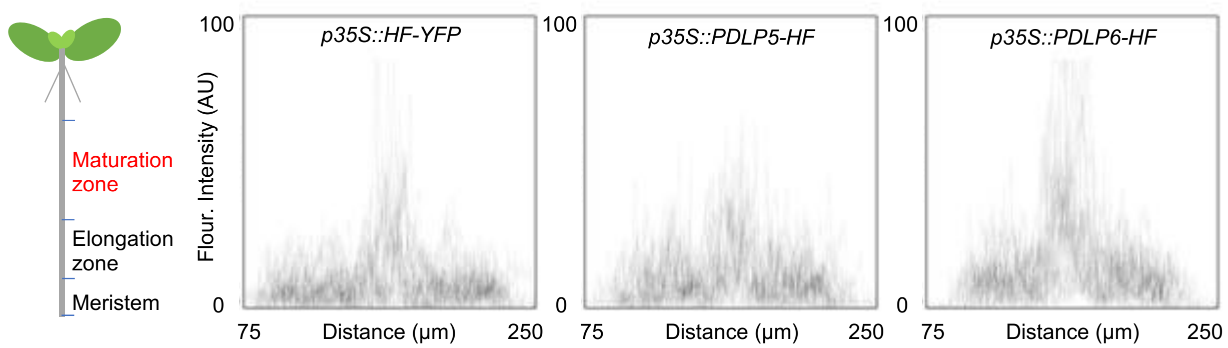


**Supplemental Figure 4. Callose accumulation in roots of Arabidopsis transgenic plants overexpressing HF-YFP, PDLP5-HF, or PDLP6-HF.** Semi-quantitative evaluation of the relative level of aniline blue-stained callose in root cells was performed by analyzing the signal intensity across different root cell types. Confocal images were captured from the maturation zone of 10-day old seedlings. 25 individual transgenic plants were analyzed for each genotype and fluorescence intensity profiles (arbitrary unit: AU) of callose accumulation were combined within the genotype. Numbers on the X axis indicate distance (μm) across the region analyzed.

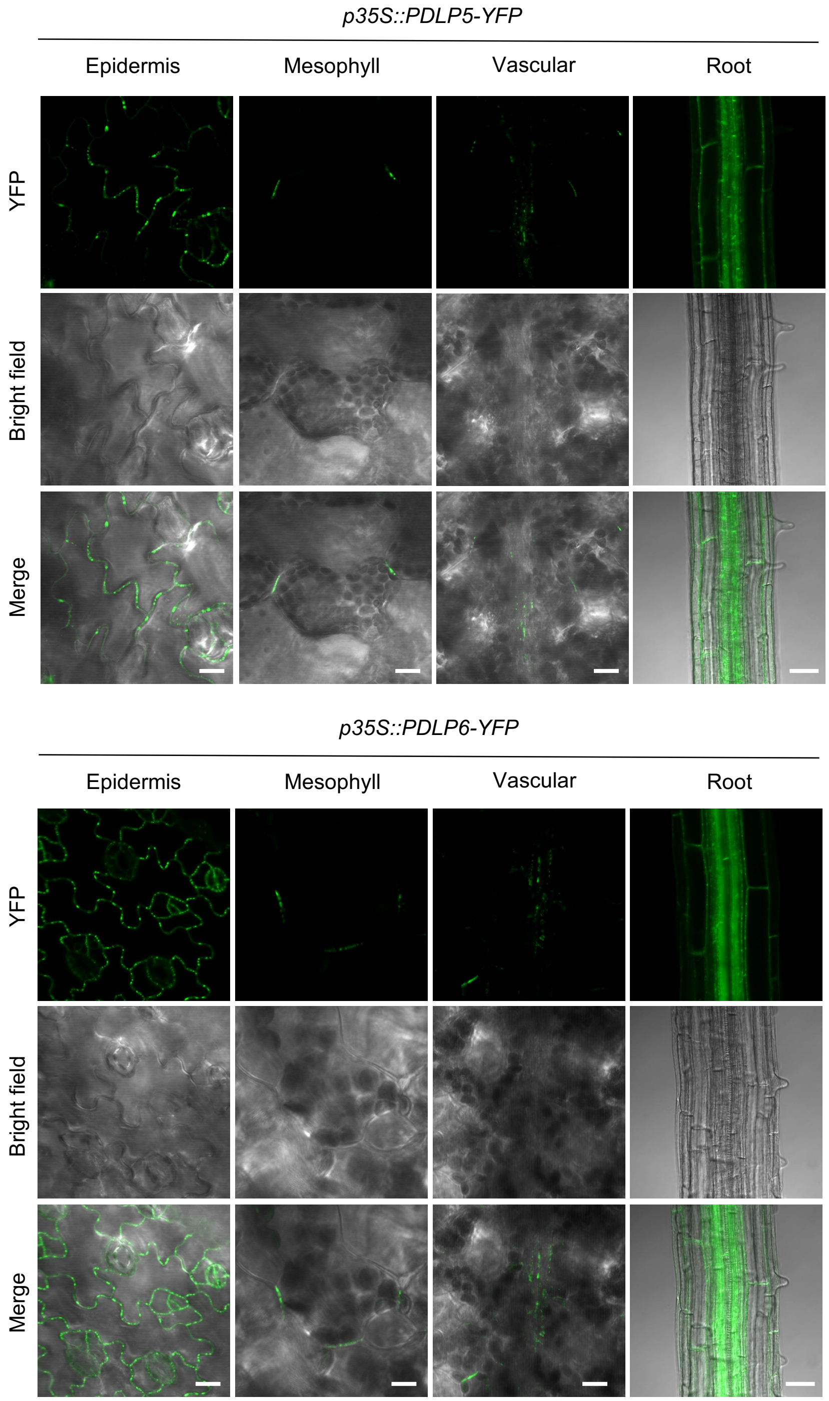


**Supplemental Figure 5. Ubiquitous expression of PDLP5-YFP and PDLP6-YFP fusion proteins in *p35S::PDLP5-YFP* and *p35S::PDLP6-YFP* transgenic plants.** The expression of the fusion proteins was detected in epidermal cells, mesophyll cells, vascular cells in leaves, and most cell types in roots. Confocal images were captured from 2-week-old Arabidopsis seedlings. Scale bars for epidermis, mesophyll, and vasculature = 10 μm. Scale bars for root = 50 μm.
